# Estimating fish population abundance by integrating quantitative data on environmental DNA and hydrodynamic modelling

**DOI:** 10.1101/482489

**Authors:** Keiichi Fukaya, Hiroaki Murakami, Seokjin Yoon, Kenji Minami, Yutaka Osada, Satoshi Yamamoto, Reiji Masuda, Akihide Kasai, Kazushi Miyashita, Toshifumi Minamoto, Michio Kondoh

## Abstract

We propose a general framework of abundance estimation based on spatially replicated quantitative measurements of environmental DNA in which production, transport, and degradation of DNA are explicitly accounted for. Application to a Japanese jack mackerel (*Trachurus japonicus*) population in Maizuru Bay revealed that the method gives an estimate of population abundance comparable to that of a quantitative echo sounder method. These findings indicate the ability of environmental DNA to reliably reflect population abundance of aquatic macroorganisms and may offer a new avenue for population monitoring based on the fast, cost-effective, and non-invasive sampling of genetic information.

---

Knowledge on the distribution and abundance of species is crucial for ecology and related applied fields such as wildlife management and fisheries. The detection and quantification of environmental DNA (eDNA) is an emerging methodology for ecological studies and could enhance the ability of investigators to infer occurrence and abundance of species. This approach has been applied, especially but not limited to, to aquatic species such as fish and amphibians and has been identified as a powerful and yet cost-effective tool for species detection (Bohmann *et al.* 2014, Rees *et al.* 2014, Thomsen & Willerslev 2015, Goldberg *et al.* 2016, Deiner *et al.* 2017, Hansen *et al.* 2018). Challenges remain, however, in quantitative applications of eDNA. Since earlier studies revealed positive correlations between species abundance and eDNA concentration (Takahara *et al.* 2012, Thomsen *et al.* 2012, Goldberg *et al.* 2013, Pilliod *et al.* 2013, Eichmiller *et al.* 2014), it has been expected that local population abundance may be inferred by measuring the concentration of eDNA at a given locality. Indeed, an analytical framework proposed recently for eDNA-based abundance estimation assumes a probability distribution that represents the quantitative relation between eDNA concentration and the underlying population size (Chambert *et al.* 2018). Nonetheless, such a definite relation may not always be present, possibly depending on e.g. the shedding rate, transport, and exogenous input of eDNA (Pilliod *et al.* 2013, Eichmiller *et al.* 2014, Lacoursière-Roussel *et al.* 2016, Yamamoto *et al.* 2016, Jo *et al.* 2017).

The fundamental factors that underlie such context dependency are the ‘ecology of eDNA’: the distribution of eDNA in space and time stems from processes governing the origin, state, transport, and fate of eDNA particles (Barnes & Turner 2016). Thus, in applications of the eDNA methodology, detailed information about such processes may be critical. Without relevant knowledge of these processes, for example, the spatial and temporal scales of information provided by eDNA remain largely uncertain (Thomsen & Willerslev 2015, Goldberg *et al.* 2016, Hansen *et al.* 2018). Therefore, here, our purpose was to develop a general approach to eDNA-based abundance estimation that can fully account for the ecology of eDNA, i.e. the rate of production and degradation of eDNA as well as the transport of eDNA within a flow field in an aquatic area of interest. We use a *tracer model*: a numerical hydrodynamic model that can simulate the distribution of eDNA concentrations within an aquatic area. Under certain assumptions, the behaviour of the model can also be regarded mathematically as a linear function of an input vector representing the distribution of population abundance levels (densities) within the area. The inference of population abundance then boils down to Bayesian estimation of coefficients of a generalised linear model (see Methods for details).

We applied this approach to a population of the Japanese jack mackerel (*Trachurus japonicus*, a commercially important fish species) in Maizuru Bay, Japan (Fig. 1). The bay has a surface area of ∼22.87 km^2^ with a maximal water depth of approximately 30 m, where the jack mackerel is numerically the most dominant fish species. The field work was conducted during a season in which the jack mackerel population in the bay is dominated by new recruits. We cruised the bay on two days in June 2016 to collect water samples at 100 stations and to conduct an acoustic survey. On the basis of the eDNA concentration measurements and a tracer model configured for Maizuru Bay, we obtained an estimate of fish population abundance in the bay. This estimate was then verified via a parallel estimate of abundance obtained by a quantitative echo sounder method.

**Fig. 1.**
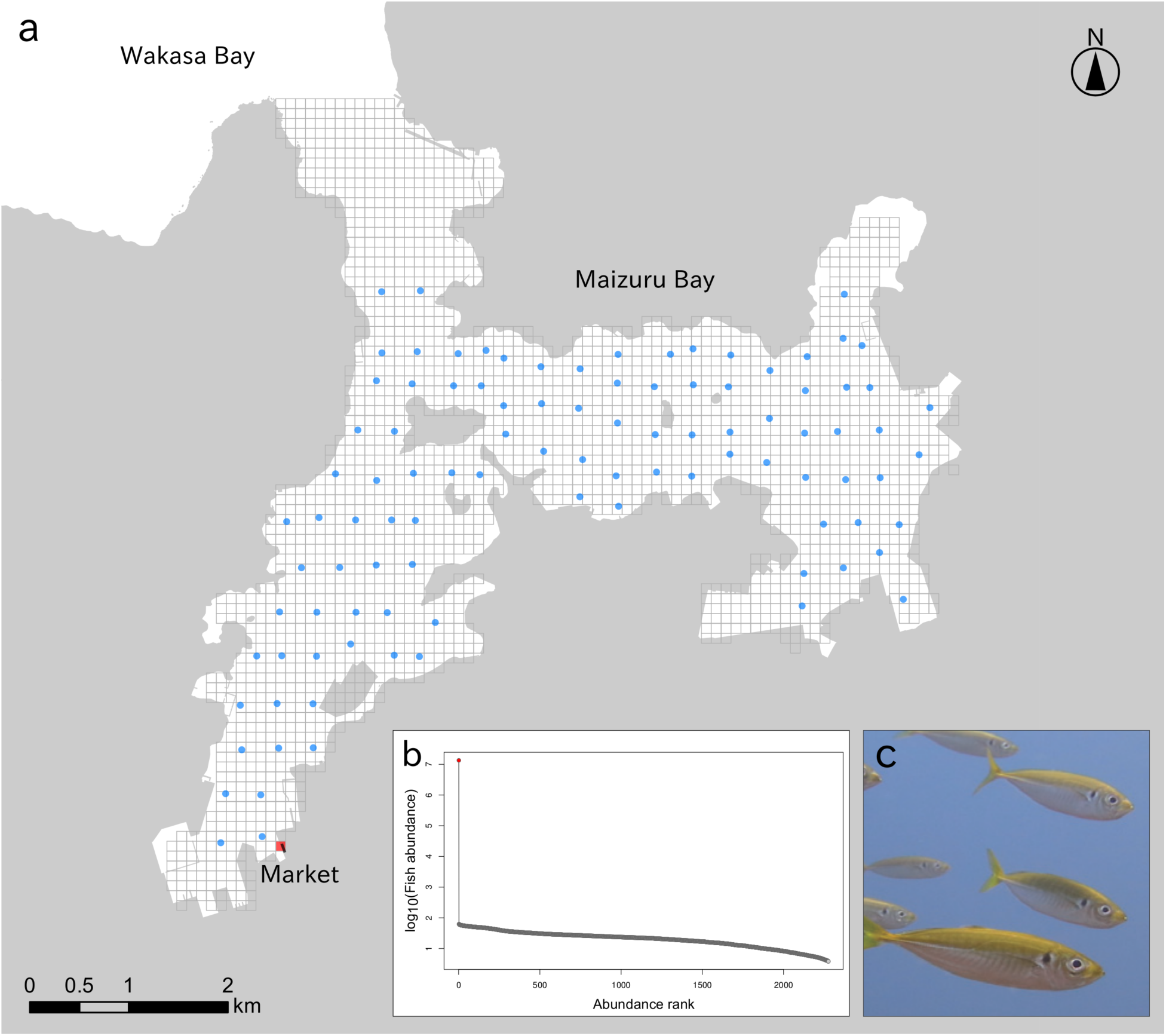
Maizuru bay, the study site. a, The 2,276 horizontal lattice grids for the eDNA tracer model (grey boxes) and the 100 water-sampling stations (blue circles). The grid in which estimates of jack mackerel density were extremely high is highlighted in red. The building of the fish market, overlapping with the red lattice grid, is depicted by a filled black box. b, Fish abundance estimates in the 2,276 horizontal lattice grids. Abundance estimates in nine vertical cells were pooled for each grid. The lattice grid next to the market is highlighted in red. c, The Japanese jack mackerel (*T. japonicus*) in Maizuru bay (photo credit: R. Masuda).

The abundance estimates yielded by the two methods were comparable; the point estimate of the eDNA method was of the same order of magnitude as that of the quantitative echo sounder estimate, which was covered by the 95% highest posterior density interval (HPDI) of the eDNA-based estimate (Table 1). Moreover, we could identify a coordinate of grids in which density of jack mackerels was estimated to be unrealistically high; fish abundance in this location was estimated at as much as tens of millions of individuals (posterior median and 95% HPDI: 1.35 × 10^7^ [0.00 to 1.77 × 10^7^] individuals; Fig. 1b). It is located next to a wholesale fish market (Fig. 1a), which has been suspected as a significant source of exogenous jack mackerel eDNA in Maizuru Bay (Yamamoto *et al.* 2016, Jo *et al.* 2017). We therefore regarded the extreme estimates in these cells as resulting from a massive eDNA input from the market and excluded them from the inference of the bay scale fish abundance. This correction reduced the estimate of fish abundance in the bay, whereas the 95% HPDI still covered the echo sounder estimate (Table 1).

**Table 1.** Estimates of Japanese jack mackerel abundance in Maizuru Bay. The second row of the eDNA method gives the abundance estimate that excluded the grid cells close to the wholesale fish market (indicated in Fig. 1a), which were identified as extraordinary eDNA sources. The point abundance estimates and credible intervals are presented as posterior medians and highest posterior density intervals, respectively. In both estimation methods, estimates are obtained under the assumption that the size of jack mackerel individuals was 3 cm in body length and 1 g in body weight (see Methods).

| Method | Abundance estimates | 95% Bayesian credible interval |
| --- | --- | --- |
| eDNA + hydrodynamic model | $3.31 \times 10^7$ | $(2.32 \times 10^7, 6.32 \times 10^7)$ |
| (fish market cells omitted) | $2.23 \times 10^7$ | $(0.77 \times 10^7, 5.29 \times 10^7)$ |
| Quantitative echo sounder | $3.91 \times 10^7$ | — |

The eDNA methods are rapidly developing technologies that have a great potential to facilitate the understanding and management of aquatic species, although their quantitative applications are still the critical step. A number of quantitative eDNA applications uncovered a positive association between eDNA concentration and abundance of a target species (Takahara *et al.* 2012, Thomsen *et al.* 2012, Goldberg *et al.* 2013, Pilliod *et al.* 2013, Eichmiller *et al.* 2014, Lacoursière-Roussel *et al.* 2016, Yamamoto *et al.* 2016, Jo *et al.* 2017). With the aid of a well-designed sampling scheme and an associated statistical model, such relations can help to quantify abundance at multiple locations, especially in lentic systems where advection of eDNA is limited (Chambert *et al.* 2018). This study presents a novel approach to abundance estimation based on quantitative eDNA measurements into which a numerical hydrodynamic model (i.e. the tracer model) is incorporated to explicitly account for the details of the ecology of eDNA. It may be flexibly applied to a wide array of aquatic systems in which hydrodynamics and rates of eDNA shedding and degradation are modelled, thereby broadening the scope of the general idea implemented recently in a one-dimensional lotic system with a single eDNA source (Sansom & Sassoubre 2017) and in a river network system with multiple eDNA sources (Carraro *et al.* 2018). The application of the proposed approach to the Japanese jack mackerel population in Maizuru Bay indicates that abundance of species can be reliably estimated by means of eDNA in a mesoscale lotic system. Furthermore, the results revealed that the method can distinguish major exogenous sources of eDNA, which have been recognised as a nuisance factor in eDNA applications especially for species subject to fishery (Yamamoto *et al.* 2016, Jo *et al.* 2017).

The proposed framework, however, has several limitations in its current form. It requires several key assumptions, such as the stationarity (i.e. demographic closure) of the population and homogeneity of individuals in terms of their rate of eDNA shedding. In addition, the number of eDNA samples may typically be smaller than the number of grid cells in the tracer model, thus requiring some sort of models explaining the association between population density and measured covariates and/or regularisation (i.e. prior specification) to make a statistical inference (see Methods). Although our results indicated that the method can be applied even with these limitations, further methodological development would be warranted. A promising approach among quantitative eDNA applications is to combine eDNA measurements and classical protocols for abundance estimation (Chambert *et al.* 2018); this strategy is also likely to improve the general approach proposed here.

It has been argued that in an application of the eDNA method, careful consideration of details of the ecology of eDNA is critical (Bohmann *et al.* 2014, Rees *et al.* 2014, Thomsen & Willerslev 2015, Barnes & Turner 2016, Goldberg *et al.* 2016, Deiner *et al.* 2017, Hansen *et al.* 2018). We implemented this idea in a quantitative eDNA method, leading to integration of eDNA concentration measurements and hydrodynamic modelling for abundance estimation. Because the research on aquatic eDNA of macroorganisms is still in its infancy since its discovery (Ficetola *et al.* 2008), more work is needed to elucidate the processes that determine a distribution of eDNA in the field; knowledges on the ecology of eDNA will help to improve the accuracy of quantitative eDNA approaches. The relatively less explored field of quantitative eDNA applications lies in the multispecies context, which involves eDNA metabarcoding rather than the targeted quantitative PCR (qPCR) method (Deiner *et al.* 2017). A quantitative metabarcoding technique (Ushio *et al.* 2018) may hold great promise for enabling researchers to analyse many aquatic species at a time. Exploring between-species differences in the rate of eDNA shedding and degradation may therefore be worthwhile. In addition to remarkable efficiency in species detection, we expect that eDNA methodologies can enhance the ability of investigators to gain quantitative insights into aquatic ecosystems.

## Methods

### A general framework for abundance estimation

#### The tracer model as a linear projection function

Here, we define a *tracer model* as a numerical hydrodynamic model that simulates generation, transport, and decay of particles (i.e. eDNA) on the basis of a flow field determined by given physical conditions within an aquatic area of interest. In this study, we assume a tracer model for a three-dimensional discrete space in which the entire aquatic area of interest is discretised into grid cells of known volume. A tracer model can in principle simulate the ecology of eDNA and thus derives a spatial distribution of eDNA within the aquatic area, given that per capita and unit time shedding rates of eDNA, degradation rates of eDNA, and density (or equivalently, abundance) of organisms in each grid cell are specified, in addition to the flow field. The main idea that underlies the framework we propose is that we can regard a tracer model as a function that takes a vector of cell level density of organisms as an input and outputs eDNA concentration in each grid cell at a point in time; thus, the inference of abundance is an inverse problem: finding an input vector of a tracer model (i.e. density of organisms in each grid cell) that best explains measurements of eDNA concentration that are collected at a point in time and are replicated spatially within the aquatic area of interest.

Nevertheless, such a problem is difficult to solve under the general conditions where both the environment and abundance vary in a complex manner. We therefore make several key assumptions that simplify the problem. Firstly, we assume that during two time points *t* and *s* (< *t*), key environmental variables for hydrodynamic processes are known from some observations and/or model prediction so that the flow field can be determined and plugged in to the tracer model. Here, *t* refers to the point in time at which eDNA concentration is observed at multiple locations within the aquatic area, and *s* denotes some point in time sufficiently far away from *t* such that eDNA concentration at *t* is virtually independent from that at *s*. Secondly, we assume that the rates of production and degradation of eDNA are known in each grid cell during the period between *s* and *t*. They may either be regarded as constant across space and time or assumed to vary depending on known environmental variables, such as water temperature, salinity, and pH, so that the rates of generation and disappearance of eDNA can be determined completely in the tracer model. In addition, we assume that these rates are independent of the eDNA concentration, and thus both production and degradation of eDNA are linear processes. Third, we suppose that in each grid cell, all eDNA particles arise exclusively from individuals of the target species that are identical in their eDNA-shedding rate. Finally, we assume that abundance is stationary in each grid cell throughout the period between *s* and *t* (i.e. the demographic closure assumption; Williams *et al.* 2002).

Under these assumptions, a tracer model can be regarded as a linear function. We denote density of organisms in cell *i* (*i* = 1, …, *M*) by *x*_*i*_ and define **x** = (*x*_1_, *x*_2_, …, *x_M_*). Let us denote the water volume of each cell by **v** = (*v*_1_, *v*_2_, …, *v_M_*) so that abundance in mesh *i* and in the whole aquatic area is expressed as *v_i_x_i_* and **v**^*T*^**x**, respectively (here, **a**^*T*^ means the transpose of vector **a**). The tracer model predicts eDNA concentration in each grid cell at time point *t* that results from the generation, advection, diffusion, and degradation of eDNA occurring between *s* and *t* within a given flow field, which we denote (without an explicit index of *t*) by **w** = (*w*_1_, *w*_2_, …, *w_M_*). If *a*_*ij*_ is defined as the (per unit density) contribution of mesh *j* to eDNA concentration in mesh *i* at time *t*, then eDNA concentration can be expressed as *w*_*i*_ = *a*_*i*1_*x*_1_ + *a*_*i*2_*x*_2_ + *· · ·* + *a_iM_ x_M_*. If we designate **A** = (*a*_*ij*_)_*M×M*_, then this equation can be written in a matrix form as **w** = **Ax**. Thus, although a tracer model indeed represents temporal evolution of eDNA concentration within the period between *s* and *t* according to some differential equations (presented below), its behaviour can be described simply — under the assumptions noted above — by matrix **A**, which projects the vector of density **x** onto the vector of eDNA concentration **w**. For *i* = 1, …, *M*, the *i*th column of **A** can be obtained numerically as a result of execution of the tracer model between time points *s* and *t* with a vector of density in which cell *i* has a unit density and all other cells have 0 density.

#### Fitting the tracer model to eDNA concentration data

We assume that eDNA concentration was measured in *N* samples collected within the aquatic area of interest at a point in time (or, in practice, within a sufficiently short period). Let us denote the observed eDNA concentration in sample *n* by *y*_*n*_ (*n* = 1, …, *N*) and express it with vector **y** = (*y*_1_, …, *y_N_*). In the following text, we suppose that all eDNA measurements are positive (i.e., *y*_*n*_ > 0). Note, however, that negative samples could also be included in the analysis given that the detection process of eDNA is modelled jointly (Carraro *et al.* 2018). We define *i*(*n*) as an index variable that means the index of the cell in which sample *n* was obtained. If we let **B** = (*ai*(*n*)*j*)_*N×M*_, the prediction of the tracer model for the data vector, as a function of density vector **x** is then expressed as **Bx**.

Because the tracer model yields a linear predictor for **y**, we can apply the (generalised) linear modelling approach (McCullagh & Nelder 1989) to estimate density vector **x**; in particular, we can regard **B** and **x** as a design matrix and a vector of coefficients of a linear regression model, respectively (note that because **x** represents density, the searches for estimates should be within the space of parameters such that *x_i_* ≥ 0 for all *i*). For example, we can consider the following normal linear model:

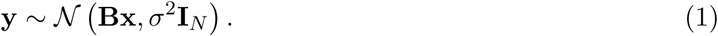

where 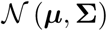 is a multivariate normal distribution with mean vector ***µ*** and covariance matrix **Σ**, *σ*^2^ is a residual variance of the linear model, and **I**_*m*_ is a *m × m* identity matrix. A maximum likelihood estimation gives estimate 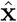 that minimises the residual square error 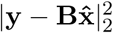. Alternatively, we can fit the model on a logarithmic scale; this approach may be more reasonable than the above model when a lognormal error structure better represents eDNA concentration data as is often the case in quantitative eDNA studies (e.g. Takahara *et al.* 2012, Thomsen *et al.* 2012, Eichmiller *et al.* 2014, Wilcox *et al.* 2016, Jo *et al.* 2017). The alternative model can be written as

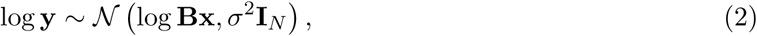

which is a generalised linear model with an exponential link function: a less popular but still appropriate within the generalised linear modelling framework given that *x_i_* ≥ 0 for all *i* (McCullagh & Nelder 1989). The maximum likelihood method for this model yields estimate **x**ˆ that minimises the residual square error 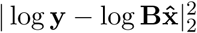.

The standard maximum likelihood approach is, however, not applicable to these models when

*M > N* because the maximum likelihood estimate of **x** is not uniquely identified in this setting. This may be a typical situation at a reasonable level of spatial discretisation for the tracer model and sampling effort of eDNA. When some covariates, assumed to covary with density, are available for each cell, a (generalised) linear model for density can be introduced to effectively reduce the number of unknown parameters (Carraro *et al.* 2018). Specifically, density of the target species can be modelled, for example, as log **x** = **Z*β***, where **Z** is a matrix of covariates, and ***β*** is a vector of coefficients (including an intercept). Otherwise, additional regularisation is necessary to make an inference based on such singular models. The regularisation method often employed for regression models is to impose a penalty on the size of regression coefficients; a typical example includes ridge regression and lasso, which can be interpreted in general as a Bayesian inference of the model with a specific prior on the regression coefficients (Hastie *et al.* 2009). Thus, inference can be achieved via a Bayesian model-fitting approach such as empirical Bayes and the full-Bayesian inference (Karabatsos 2018).

### An application to a marine fish population

#### The Japanese jack mackerel in Maizuru Bay

The study was conducted in Maizuru Bay (Kyoto prefecture, Japan; 35 29′ N, 135 23′ E) to estimate abundance of the jack mackerel (*T. japonicus*) via concentration of eDNA. The bay has a surface area of ∼22.87 km^2^ with a maximum water depth of approximately 30 m, and connects with Wakasa Bay through a narrow bay mouth in its north (Fig. 1).

According to long-term underwater visual surveys, the jack mackerel is numerically the most dominant fish species in shallow (< 10 m in depth) coastal waters in this area (Masuda 2008); their body size ranges from 10 to 45 mm in standard length offshore and 40–120 mm standard length in the shallow rocky reef habitat (Masuda *et al.* 2008). The study was conducted during the peak season of jack mackerel recruitment from the offshore pelagic zone to a coastal shallow reef habitat, where the jack mackerel population in the bay is dominated by new recruits. In the following analysis, we therefore assumed that the population is represented by individuals of size ∼ 3 cm (body length) and ∼ 1 g (body weight; see Supplementary information).

#### Measurement of eDNA concentration

We conducted the water sampling on 21 and 22 June 2016 from a research vessel at 100 stations located approximately on ∼400 m grids in Maizuru Bay (Fig. 1). Samples were collected at 53 stations in the eastern part of the bay on the first day and at 47 stations in the western part on the second day. The average water depth at the 100 stations was ∼15m. On both days, the survey began from the mouth of the bay and ended in the inner most part of the bay. The survey was approved by the harbourmaster of Maizuru Bay (Permission number 160 issued on 5th May 2016).

At each sampling station, we captured sea water at three depths: the surface, middle, and bottom. The middle and bottom depths were defined as 5 m from the surface, which was just below the pycnocline, and 1 m above the sea floor, respectively. Water samples were collected with a ladle for surface water and vanDorn samplers for middle and bottom water. For each station and depth, a 1 L water sample was placed in a plastic bottle, which was rinsed in advance with a subset of captured water. We then immediately added 1 mL of 10% benzalkonium chloride to the samples and mixed them gently to prevent degradation of DNA (Yamanaka *et al.* 2017). The bottles of water samples were stored in opaque containers to avoid sunlight.

We filtered water samples on the same day of the field survey through a 47 mm diameter glass microfiber filter (nominal pore size 0.7 *µ*m, GE Healthcare Life Science [Whatman]) using an aspirator in a laboratory at Maizuru Fisheries Research Station, Kyoto University. The filters were folded so that the filter surface faced inward and were wrapped into aluminium foil to store at −20 C until eDNA extraction. It took less than 7 h to complete all operations from the water collection to the filtration. To prevent carryover of eDNA, filtration devices were bleached by means of 0.1% sodium hypochlorite for at least 5 min and then were washed and rinsed with tap water and distilled water, respectively, to clear the remaining sodium hypochlorite. This bleaching process was validated by a series of negative controls of filtration undertaken for every sequence of 15 filtrations in which 1 L of distilled water was filtered with bleached equipment.

All samples and negative controls of filtration were subjected to eDNA extraction and subsequent quantitative PCR (qPCR). eDNA extraction was conducted by following the procedure of Yamamoto *et al.* (2016), which eventually yielded 100 *µ*L of a DNA solution. We determined the concentration of mitochondrial cytochrome b (CytB) of the jack mackerel by qPCR on a LightCycler 96 machine (Roche). The primers and probe used in the qPCR were as follows: forward primer, 5′–CAG ATA TCG CAA CCG CCT TT–3′; reverse primer, 5′–CCG ATG TGA AGG TAA ATG CAA A–3′; probe, 5′–FAM-TAT GCA CGC CAA CGG CGC CT–TAMRA–3′ (Yamamoto *et al.* 2016). This primer set amplifies 127 bp of the CytB gene. The PCR reaction solution was 20 *µ*L: 2 *µ*L of the extracted DNA solution, a final concentration of 900 nM forward and reverse primers and 125 nM TaqMan probe in 1 × TaqMan master mix (TaqMan gene expression master mix; Life Technologies). The thermal program for the qPCR was as follows: 2 min at 50°C, 10 min at 95 C, and 55 cycles of 15 sec at 95°C and 1 min at 60°C. To draw quantification standard curves, we simultaneously performed PCR on 2 *µ*L of artificial DNA solutions that contained 3 × 10^1^ to 3 × 10^4^ copies of our target sequence. qPCR was carried out in triplicate for each sample and standard. In addition, a 2 *µ*L pure water sample was analysed simultaneously in triplicate as a negative control of PCR. In all the runs, *R*^2^ values of calibration curves were more than 0.99, the range of slopes was between −3.859 and −3.512, and the range of *y*-intercepts was between 38.34 and 40.36. No eDNA of the jack mackerel was detected in any negative control sample of filtration and PCR.

#### Development of the tracer model

To obtain the flow field in Maizuru Bay, we configured the Princeton ocean model (POM) with a scaled vertical coordinate (i.e. the sigma coordinate system; Mellor 2002) for the bay. The model represented Maizuru Bay by 20,484 grid cells. Specifically, the bay was discretised by 2,276 horizontal lattice grids at a resolution of 100 m, and the grids had nine non uniform vertical layers, with finer resolution near the surface; the sigma coordinate was set as *σ* = 0.000, −0.041, −0.088, −0.150, −0.245, −0.374, −0.510, −0.646, −0.796, and −1.000. The configuration of the model was achieved by means of the bottom topography of the bay, data and model estimates of surface meteorological conditions, estimated river discharges, and the model results of Wakasa Bay as the open boundary conditions (Yoon & Kasai 2017); additional details are described elsewhere (Kasai & Yoon 2018). The model simulated flow fields within the bay from 1 June 2016, under the initial conditions interpolated from the model results of Yoon & Kasai (2017), to the final day for the water sampling (i.e. 22 June 2016). Time steps of the simulation were set to 0.1 s for the external mode and 3 s for the internal mode.

We then incorporated eDNA of jack mackerels into the POM configured for Maizuru Bay as a passive tracer to simulate its concentration within the flow field. The evolution of eDNA concentration in a grid cell, denoted by *c*, is represented as

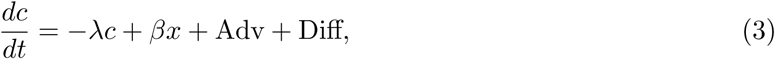

where *x* is the density of jack mackerels in the cell, *λ* represents a degradation rate of eDNA, and *β* is a per-capita shedding rate of jack mackerel DNA. Adv and Diff are the advection and diffusion terms, respectively, which were determined implicitly by running the POM for Maizuru Bay. The eDNA degradation rate was assumed to be constant and was adopted from an estimate obtained in tank experiments where the same species-specific primer set was employed (*λ* = 0.044 h^−1^; Jo *et al.* 2017). The eDNA shedding rate of the jack mackerel was assumed to be constant; it was derived mathematically and found to be *β* = 9.88 × 10^4^ copies per individual per hour, according to the results of tank experiments conducted by Maruyama *et al.* (2014) and Jo *et al.* (2017). Details of this derivation are provided in Supplementary information.

#### Estimation of jack mackerel abundance based on eDNA and the tracer model

We fitted the logarithmic model (Eq. 2) to the eDNA concentration data collected in Maizuru Bay. During the model fitting, we omitted negative samples in which the number of remaining observations was *N* = 729. For vector of density **x**, we specified an independent lognormal prior with unknown prior mean *µ* and standard deviation *τ*:

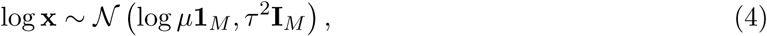

where **1**_*M*_ represents a vector of all ones with length *M*. Because *N* was significantly smaller than

*M*, we were pessimistic about estimating the spatial variation in cell level density with reasonable precision. Our main goal of the inference was therefore to quantify the bay level abundance **v**^*T*^**x** along with its uncertainty.

With uniform positive priors on *µ*, *τ*, and *σ*, we fitted the model via a fully Bayesian approach. Posterior samples were obtained by the Markov chain Monte Carlo (MCMC) method implemented in Stan (version 2.18.1; Carpenter *et al.* 2017) in which three independent chains of 10,000 iterations were generated after 1,000 warm-up iterations. Each chain was thinned at intervals of 10 to save the posterior sample.

Convergence of the posterior was checked for each parameter with the 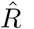 statistic. Posterior convergence was achieved at a recommended degree (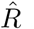 < 1.1; Gelman *et al.* 2013) in almost all parameters except log *x* in four cells. We decided, however, that the results are solid because the posterior of the bay level abundance — the target of the inference — fully converged. The goodness-of-fit assessment of the model, measured by the *χ*^2^-discrepancy statistic (Conn *et al.* 2018), gave no clear indication of a lack of model fit (Bayesian *p* value: 0.404).

#### Estimation of jack mackerel abundance from quantitative echo sounder data

An independent estimate of jack mackerel abundance was obtained based on a calibrated quantitative echo sounder by a standard acoustic survey method (Simmonds & MacLennan 2005). The acoustic survey was conducted during the survey cruise for the water sampling (described above). We used the KSE300 echo sounder (Sonic Co. Ltd., Tokyo, Japan) with two transducers (T-182, 120 kHz, and T-178, 38 kHz; beam type, split-beam; beam width, 8.5°; pulse duration, 0.3 ms; ping rate, 0.2 s), which were mounted off the side of the research vessel at a depth of 1 m. The acoustic devices were operated during the entire survey cruise to record all acoustic reflections, except when the research vessel stopped at each sampling station where the recording was stopped to avoid reflection from the sampling gear and cables. The research vessel ran at ∼4 knots, on average, between the sampling stations. The echo intensity data were denoised and cleaned in Echoview ver. 9.0 (Echoview Software Pty. Ltd., Tasmania, Australia). We omitted signals between the sea bottom and 0.5 m above it to exclude the acoustic reflection from the sea floor. Additionally, we eliminated signals from sea nettles (*Chrysaora pacifica*) by filtering reflections of −75 dB.

From the obtained acoustic data, the reflections of jack mackerel were extracted by the volume back scattering strength difference (Δ*S*_*V*_) method (Miyashita *et al.* 2004, Simmonds & MacLennan 2005). Δ*S*_*V*_ was defined as the difference in the volume backscattering strength (*S*_*V*_) between the two frequencies as follows:

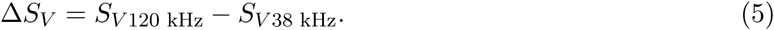

According to field validation in Maizuru Bay combining acoustic surveys and visual confirmation of jack mackerel schools by snorkelling, we assumed the range of Δ*S*_*V*_ of jack mackerel between −6.4 and 5.2 dB. This criterion discriminates the jack mackerel from larval Japanese anchovy (*Engraulis japonicus*), the subdominant species in the bay (Masuda 2008), which reflects the high frequency echo strongly as compared to low frequency (Ito *et al.* 2011) and was used to determine *S*_*V*_ of the jack mackerel in 1 m^3^ water cubes in Echoview ver. 9.0.

Density of jack mackerel in a 1 m^3^ water cube, denoted by *D*, was estimated as

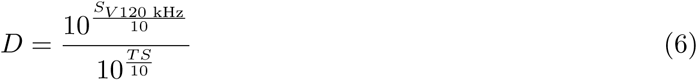

where *TS* is the target strength of an individual jack mackerel. By assuming that jack mackerel population in the bay was dominated by individuals of the size 3 cm, we chose *TS* = −59.6 dB (Nakamura *et al.* 2013, Yamamoto *et al.* 2016). The fish density on the echo sounder track lines was then matched with the grid specification of the tracer model by a box averaging method. In particular, fish density in each grid cell was estimated by a geometric mean of *D* taken over a 500 m square block that surrounds the grid cell. For grid cells in which any *D* was not available in their square block owing to a lack of the acoustic data, fish density was estimated by means of a geometric mean of the fish density taken across the other grid cells. Finally, the bay level abundance was estimated as a sum of the product of grid level density and water volume of each cell.

## Data availability

The datasets generated and analysed during the current study are available from the corresponding author upon reasonable request.

## Acknowledgements

We are grateful to the following people for the assistance with and support of the fieldwork and laboratory experiments: K. W. Suzuki, H. Sawada, Y. Ogura, M. Mukai, M. Yamashita, M. Shiomi, M. Ogata, T. Yoden, A. Fujiwara, S. Hidaka, T. Jo, M. Sakata, S. Tomita, Q. Wu, S. Ikeda, M. Hongo, S. Sakurai, N. Shibata, S. Tsuji, H. Yamanaka, M. Ogawa, M. Tomiyasu, H. Araki, H. Kawai and M. Miya. Funding was provided by the Japan Science and Technology Agency (CREST, grant number JPMJCR13A2). This research was supported by allocation of computing resources of the SGI ICE X and HPE SGI 8600 supercomputers from the Institute of Statistical Mathematics.

## Author contributions

M.K., R.M., A.K., K. Miyashita, and T.M. conceived and designed the eDNA survey. H.M. and S. Yamamoto conducted the molecular experiments. K. Minami and K. Miyashita analysed the echo sounder data. S. Yoon and A.K. developed the tracer model. K.F. and Y.O. designed the methodology and conducted data analyses. K.F., H.M., S. Yoon, and K. Minami led the writing of the manuscript. All the co-authors discussed the results and contributed critically to the manuscript.

## Supplementary information

### Body size of the Japanese jack mackerel

On 23rd June 2016, the day following the survey cruise, a trawl net (79 cm in diameter, 2.4 m long, 5 mm mesh size) was towed horizontally at four locations in Maizuru Bay where strong sonar signals were detected. The size of the collected jack mackerels (*n* = 6) was 35.3 *±* 2.5 mm (mean *±* SD) in standard length and 0.77 *±* 0.13 g in body weight. Underwater observation was also conducted on the same day, where approximately 50 individuals of jack mackerel juveniles, of the size 20–30 mm in body length, were found during a 10 min observation period. They were all associating with sea nettles (*Chrysaora pacifica*) either singly or by forming a small group. Given that the study was conducted during the peak season of jack mackerel recruitment from an offshore pelagic zone to a coastal shallow reef habitat where the jack mackerel population in the bay is dominated by new recruits, we supposed that the size of the jack mackerel in the population can effectively be represented by ∼30 mm in body length and 1 g in body weight.

### Derivation of the eDNA shedding rate of the Japanese jack mackerel

The eDNA shedding rate of the jack mackerel was derived mathematically from the results of tank experiments conducted by Maruyama *et al.* (2014) and Jo *et al.* (2017).

In the study by Jo *et al.* (2017), three adult Japanese jack mackerels ca. 15 cm in total length and ca. 40 g in body weight, on average, had been kept in three 200 L tanks. Filtered seawater was injected into the tanks as inlet water at a rate of 0.9 L min^−1^. Then, the eDNA concentration in the rearing water (*c*) can be expressed as

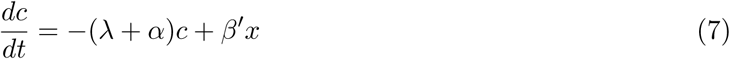

where *λ* is a degradation rate of eDNA, which had been identified in the experiment as 0.044 h^−1^ (Jo *et al.* 2017), *α* is the exponential decay constant due to water injection (0.54 h^−1^), *β*′ means the eDNA shedding rate of the adult jack mackerels, and *x* denotes the fish density in the rearing tank (0.015 individuals per litre).

We assume that the eDNA concentration had reached an equilibrium in experiments by Jo *et al.* (2017), and had been determined as *c*_0_ = 25365 copies per litre of seawater (Jo *et al.* 2017). The eDNA shedding rate of juvenile Japanese jack mackerels (*β*) is then estimated as

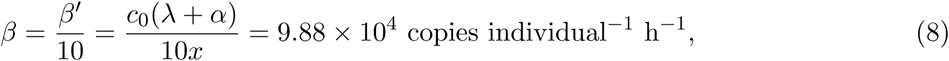

where we assumed that the eDNA shedding rate per fish body weight is four-fold higher in the juvenile fish than in the adult fish (Maruyama *et al.* 2014); this finding indicates that the eDNA shedding rate of adult individuals of size ∼ 15 cm and weight ∼ 40 g (*β*′) was 10-fold greater than that of jack mackerel juveniles of size ∼ 3 cm and weight ∼ 1 g (*β*).

